# Carbamazepine Causes Changes in Maternal Reproductive Performance and Fetal Growth Retardation in Rats

**DOI:** 10.1101/2020.09.18.303487

**Authors:** Marina Nunes, Felipe Duarte Coelho de Sousa, Rhaiza Roberta Andretta, Sandra Maria Miraglia, Samara Urban de Oliva

## Abstract

**Purpose:** Carbamazepine (CBZ) is widely used in the treatment of trigeminal neuralgia, affective disorders, and mainly as an anticonvulsant, specially by fertile women, due to their need to continuously use CBZ during pregnancy and the lactation period. CBZ crosses the placenta barrier and may impair pregnancy and the embryonic development. The aim of this study was to determine the effect of CBZ on maternal reproductive outcome, besides fetal growth and development in Wistar rats.

**Methods:** Rat dams received CBZ (20mg/Kg/day) or propylene glycol (vehicle) via intraperitoneal (i.p.) injection throughout the gestational period. On the 19^th^ day of gestation, the ovary and uterine contents were examined, and the placenta and fetuses were analyzed.

**Results:** The CBZ exposure during pregnancy caused a reduction in fetal weight, fetal weight classification, and crown-rump distance. CBZ also decreased the implantation index, average number of corpora lutea, fetal weights and crown-rump length and increased the pre and post-implantation loss rate. The CBZ-exposed fetus also presented external congenital malformations.

**Conclusion:** The results suggest that maternal exposure to CBZ interfered on several maternal reproductive outcomes and can cause severe fetal intrauterine growth restriction (IUGR).

## INTRODUCTION

The gestational period is accompanied by several physiological and anatomical changes, including hormonal oscillations, weight gain, increased plasma volume and decreased plasma proteins, These changes have significant consequences on the pharmacokinetic and pharmacodynamic properties of different therapeutic agents. Increased bioavailability and/or hepatic metabolism induction of medicinal products may occur. Some pregnant women have pre-existing medical conditions, that can compromise fetal development, either by the symptoms themselves or by exposure to potentially teratogenic drugs used in their treatment (Constantine 2014).

Epilepsy is one of the most common neurological disorders that requires continuous treatment during pregnancy. Epileptic women have a 33% risk of increased seizures and two times increase risk of hemorrhage, eclampsia, abruptio placentae, premature labor, miscarriage, stillbirth, developmental delay and major malformations. Antiepileptic drugs (AEDs) used as a monotherapy reduces the risk of adverse outcomes (Holmes 2002; Voinescu and Pennell 2015). The occurrence of an epileptic seizure in pregnant women may cause depression of the fetal heart rate and hypoxia (Pennel 2013; Yerby 1994). Pharmacological intervention in the treatment of epilepsy should be continued throughout the gestational period to reduce the risk of convulsions that are harmful to both mother and fetus.

Carbamazepine (CBZ) is an AED and mood-stabilizing drug used in the treatment of epilepsy, bipolar disorder, trigeminal neuralgia, and chronical pain disorder. This drug is widely used by women of childbearing age with epilepsy and it is considered less toxic than, phenobarbital, phenytoin, primidone or valproic acid (Fritz et al. 1976; Matalon et al. 2002).

CBZ is a lipid-soluble compound that can easily cross the placenta barrier and other biological membranes (Artama et al. 2005; Matalon et al. 2002). However, there is some controversy concerning the teratogenic effects of CBZ, various clinical studies reported malformations such as craniofacial defects, heart defects, and neural tube defects, as well as growth retardation and developmental delay. Adherence of extra-embryonic membranes and circulatory changes in the yolk sac were also observed after in vitro exposure to CBZ (Matalon et al. 2002; Piersma et al. 1998). In addition, epidemiological studies show a higher incidence of spontaneous abortions in pregnant women taking CBZ (Bech et al. 2014).

In fact, several clinical studies reported the occurrence of congenital malformations after maternal exposure to CBZ. However, few experimental studies have been conducted in order to evaluate maternal reproductive parameters and detailed analysis of the CBZ effects on the embryofetal development. The utilization and comparison between different species used as experimental models is fundamental for a better understanding of the phenomena involved embryonic and postnatal development, under different conditions. In this context, however, the rat is a useful model, since there is strong evidence that, among rodents, this species is most closely related to man (Zhao et al. 2004). Thus, the purpose of this study was to evaluate reproductive maternal outcome and embryofetal growth in rats treated with CBZ throughout pregnancy.

## MATERIAL AND METHODS

### Animals and Experimental Design

Adult female and male Wistar rats were housed in polypropylene cages (40 cm × 30 cm × 15 cm) filled with a layer of autoclaved white pine shavings, under controlled conditions: hygiene, photoperiod (12 h light/dark cycle), humidity (60%) and temperature (22-23°C). They had free access to tap water and commercial lab chow (Nuvilab, Nuvital Nutrientes). The females were mated overnight with males (two females per male); every morning, the males were separated from females and vaginal smears of each female were examined. Sperm presence in vaginal wet smears was defined as the first day of gestation (DG 1). The pregnant rats were housed individually, weighed daily and carefully monitored twice a day (morning and afternoon) for clinical signs of toxicity and possible gestational disorders.

Rat dams received CBZ (C-8981, Sigma Chemical Co., St. Louis, MO; 99.5% purity, 20mg/Kg/day; CBZ group – n=16) or propylene glycol (vehicle of CBZ; 99.8% purity; density 1.034 g/mL; 1 g/kg b.w.; Control group - C group – n=9), by via intraperitoneal (i.p.) injection, during the whole gestation. The CBZ dose chosen in the current study is the usual anticonvulsive dose used for preventing kindled seizure in Wistar rats (Cohn et al. 1978; de Feo et al. 1991; Lahtinen et al. 1996; Manent et al. 2007; McLean et al. 2004; Otsuki et al. 1998).

The i.p. injection is used in this study since it is an effectively and recognized method of drug administration in rodents, including pregnant animals; in addition, the daily administration of CBZ via gavage, during all pregnancy and lactation period, could be much more stressing to the rat dams and could interfere with the gestation period and delay the embryo development (Ahmed and El-Gareib, 2017; Manent et al. 2007). I.p. injections were given preferentially between inner thigh and extern genitals or into the lower left quadrant of the abdomen to ensure no organs were punctured (Nebendahl 2000).

### Maternal reproductive and offspring performance

On GD 19, the pregnant rats were submitted to euthanasia through CO_2_ inhalation (Boivin et al. 2017; Cartner et al. 2007). Then, laparotomy was performed and the uterus was removed and weighted. The numbers of implantations and resorptions were counted. The ovaries were also removed and the corpora lutea computed. The number of live/ dead fetuses, placenta and fetus weight and crown-rump distance were also recorded.

The fetuses were weighed and classified as: adequate (APA) - weights did not diverge more than + 1.7 standard deviations (SDs) from the control group (1.26 - 2.0 g range), small (SPA) - weights were at least 1.7 SD lower than the control group (≤ 1.25) or large (LPA) - weights were at least 1.7 SDs greater than the control group (≥ 1.99) (Damasceno et al. 2013; Soulimane-Mokhtari et al. 2005).

The fetuses were analyzed for the presence of external anomalies and sexed. The external macroscopic analysis consisted of number (absent, supernumerary, double), shape, position, size (small, enlarged, dilated, narrowed, elongated, distended, short, atresia) and color evaluation of the craniofacial structures (facial clefts, presence of exophthalmos, implantation in both ears, pinna malformations, presence and position of facial appendages, nasal deformities), observation of anterior and posterior extremities (number of fingers; presence, position and size of limbs), thoracic region, abdominal and dorsal (presence of hematoma, spina bifida, gastroschisis) and tail (presence, size and shape).

In sequence, half of the fetuses in each litter were fixed in Bodian’s fluid for visceral examination as described by Wilson (1965) and Barrow and Taylor (1969). The remaining fetuses were stored in 70% ethanol for skeletal examination after the alizarin red S staining procedure (Dawson 1926; Staples and Schnell 1964). Skeletal examination of all fetuses was made under a dissecting microscope. Anomalies such as absence, abnormal shape and size of the bones were recorded. Examination of the head, which included evaluation of the cranial, facial and palatal bones; examination of the trunk, in which anomalies of the vertebrae, ribs and sternum were recorded; examination of the bones of the forelimbs, including the bones of the metacarpus and phalanges, and examination of the bones of the hindlimbs, which, similarly, included examination of the metatarsal and phalangeal bones.

The following parameters were calculated: Implantation index [(number of implantations/ number of corpora lutea) X 100]; Pre-implantation loss rate [(number of corpora lutea - number of implantation sites /number of corpora lutea) x 100]; Post-implantation loss rate [(number of implantations - number of live fetuses / number of implantations) x 100]; Resorption rate [(number of resorption / number of implantations) x 100]; Sexual index [number of male fetus/ number of female fetus]; Placental index [placental weight/ fetal weight] (Tyl and Marr 2012).

The experimental protocol followed the ethical principles adopted by the Brazilian College of Animal Experimentation. The schedule for animal care and treatment was approved by the local Institutional Ethics Committee (Protocol number 2009/11).

### Statistical analysis

The data were submitted to statistical tests using the GraphPad Prism 8.2.1 software. The data that passed the normality test were submitted to t-test (Student’s test) and expressed as mean ± standard deviation (SD). The non-parametric Mann-Whitney’s test was used to compare data that failed the normality test and were expressed as median (interquartile range – IQR). The Fisher’s exact test was used for comparison of proportions. Differences were considered significant when p ≤ 0.05.

## RESULTS

CBZ-treated pregnant rats showed no signs of physical debility or gestational anomalies. All females survived to scheduled study termination on GD 19. However, the CBZ treatment caused a significant decrease in absolute maternal body weight from GD14 until GD19, as well as reducing weight gain in pregnant rats (Figure 1).

**Fig. 1.**
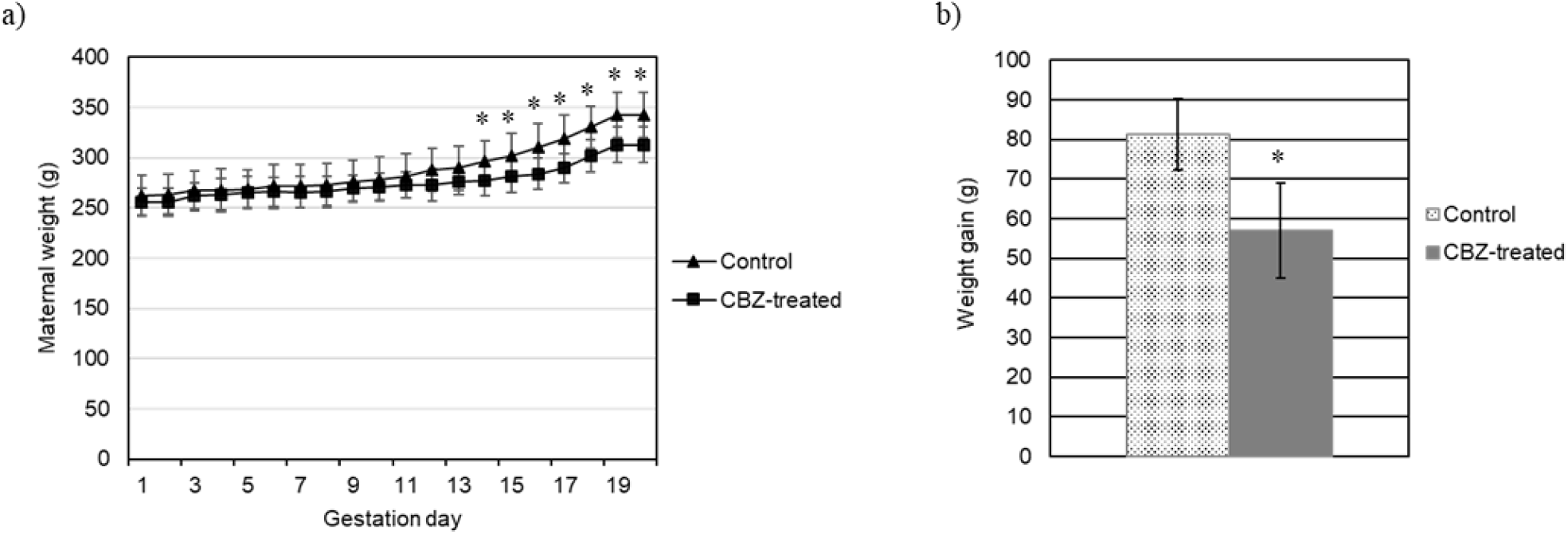
Maternal weight (A) and body total weight gain (B) of control and CBZ-treated pregnant rats during all period of pregnancy. Data are reported as mean ± SD; Student’s test. *p<0.05

The corpora lutea number, pre-implantation and post-implantation loss rate in the CBZ-treated group was greater as compared with the control group (Table 1). No significant differences were found between the CBZ-treated and the control groups for resorption and implantation numbers, post-implantation loss and resorption rates, number of pups per litter, live fetuses and sex ratio. Meanwhile, fetal body weights and fetal crown-rump lengths were smaller after prenatal CBZ exposure. Regarding placental weight and index, the CBZ-treated group did not show statistically significant differences compared to the control group (Table 1).

**Table 1.** Reproductive Outcomes from control and CBZ-treated groups.

|  | Control | CBZ -treated |
| --- | --- | --- |
| Pregnant females (n) | 9 | 10 |
| Corpora lutea |  |  |
| Total (n) | 113 | 144 |
| Median (IQR1-IQR3)<sup>B</sup> | 12 (12.0-13.0) | 14 (13.0-14.8)* |
| Implantation |  |  |
| Total (n) | 112 | 115 |
| Median (IQR1-IQR3)<sup>B</sup> | 12 (12.0-13.0) | 12.5 (11.3-13.0) |
| Implantation index (%)<sup>C</sup> | 98.9 | 81.6* |
| Total fetuses (n) | 108 | 104 |
| Number of fetuses/litter<sup>B</sup> | 12.0 (11.0-13.0) | 11.0 (9.5-12.0) |
| Live fetuses |  |  |
| Total (n) | 107 | 96 |
| Median (IQR1-IQR3)<sup>B</sup> | 12.0 (11.0-13.0) | 11 (7.5-11.8) |
| Dead fetuses |  |  |
| Total (n) | 1 | 8 |
| Median (IQR1-IQR3)<sup>B</sup> | 0.0 (0.0-0.0) | 0 (0.0-0.0) |
| Resorptions |  |  |
| Total (n) | 4 | 10 |
| Median (IQR1-IQR3)<sup>B</sup> | 0 (0.0-1.0) | 1.0 (1.0-1.0) |
| Resorption rate (%)<sup>C</sup> | 3.8 | 8.2 |
| Pre-implantation loss (%)<sup>C</sup> | 1.0 | 19.2* |
| Post-implantation loss (%)<sup>C</sup> | 4.6 | 15.6* |
| Fetal body weight (g)<sup>A</sup> |  |  |
| Mean ± SD | 1.6±0.2 | 1.4±0.2* |
| % SPA<sup>C</sup> | 2.8 | 19,2* |
| % APA<sup>C</sup> | 85.2 | 79,8* |
| % LPA<sup>C</sup> | 12.0 | 1* |
| Sex ratio (M/F)<sup>C</sup> | 1.00 | 0.87 |
| Fetal crown-rump length (mm)<sup>A</sup> |  |  |
| Mean ± SD | 24.2±1.4 | 23.2±1.1* |
| Placental weight (g)<sup>A</sup> |  |  |
| Mean ± SD | 0.42±0.05 | 0.41±0.08 |
| Placental index<sup>A</sup> |  |  |
| Mean ± SD | 0.26±0.04 | 0.29±0.05 |

IQR = interquartile range; SD = standard deviation; SPA = small for pregnancy age; APA = adequate for pregnancy age; LPA = large for pregnancy age. <sup>A</sup> Student's t-test; <sup>B</sup> Mann-Whitney test; <sup>C</sup> Fisher exact method.
\*p>0.05.

The presence of external malformations such as gastroschisis, open eyelid (ablepharia) and bilateral exophthalmos was considered related to CBZ exposure (Table 2; Figures 2 and 3). Visceral and skeletal anomalies in the head, trunk, forelimbs and hindlimbs did not differ significantly between the control and the CBZ treated groups (Table 2).

**Table 2.** Incidences of external, visceral and skeletal anomalies in fetuses from control and CBZ-treated groups.

|  | Control | CBZ-treated |
| --- | --- | --- |
| *External malformations* |  |  |
| Fetuses examined (litter) | 108 (9) | 104 (10) |
| Fetuses with anomalies (%) | 0 (0.0) | 5 (4.81) |
| Gastroschisis | 0/108 (0.0) | 1/104 (0.96) |
| Exophthalmos bilateral | 0/108 (0.0) | 1/104 (0.96) |
| Ablepharia | 0/108 (0.0) | 3/104 (2.88) |
| *Visceral malformations* |  |  |
| Fetuses examined (litter) | 54 (9) | 52 (10) |
| Fetuses with anomalies (%) | 0 (0.0) | 2 (3.85) |
| Macrophthalmia | 0 (0.0) | 1/52 (1.92) |
| Ectopic testis | 0 (0.0) | 1/52 (1.92) |
| *Skeletal malformations* |  |  |
| Fetuses examined (litter) | 54 (9) | 52 (10) |
| Fetuses with anomalies (%) | 0 (0.0) | 1 (1.92) |
| Unossified hindlimbs | 0/54 | 1/52 (1.92) |

Fisher exact test.

**Fig. 2.**
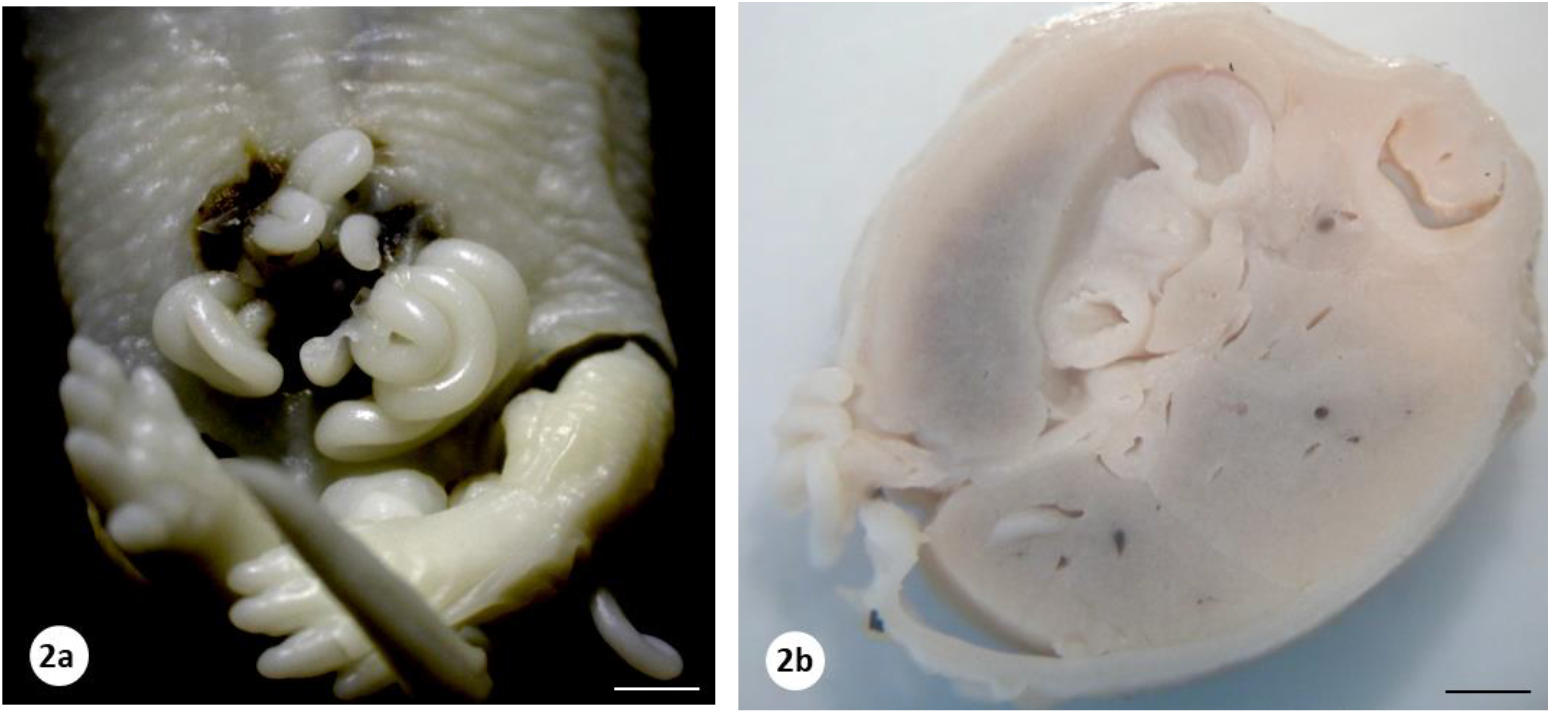
Photographs of CBZ-exposed fetus presenting gastroschisis. (a) Frontal view (b) Transversal section. Bars: 250 μm and 830μm, respectively.

**Fig. 3.**
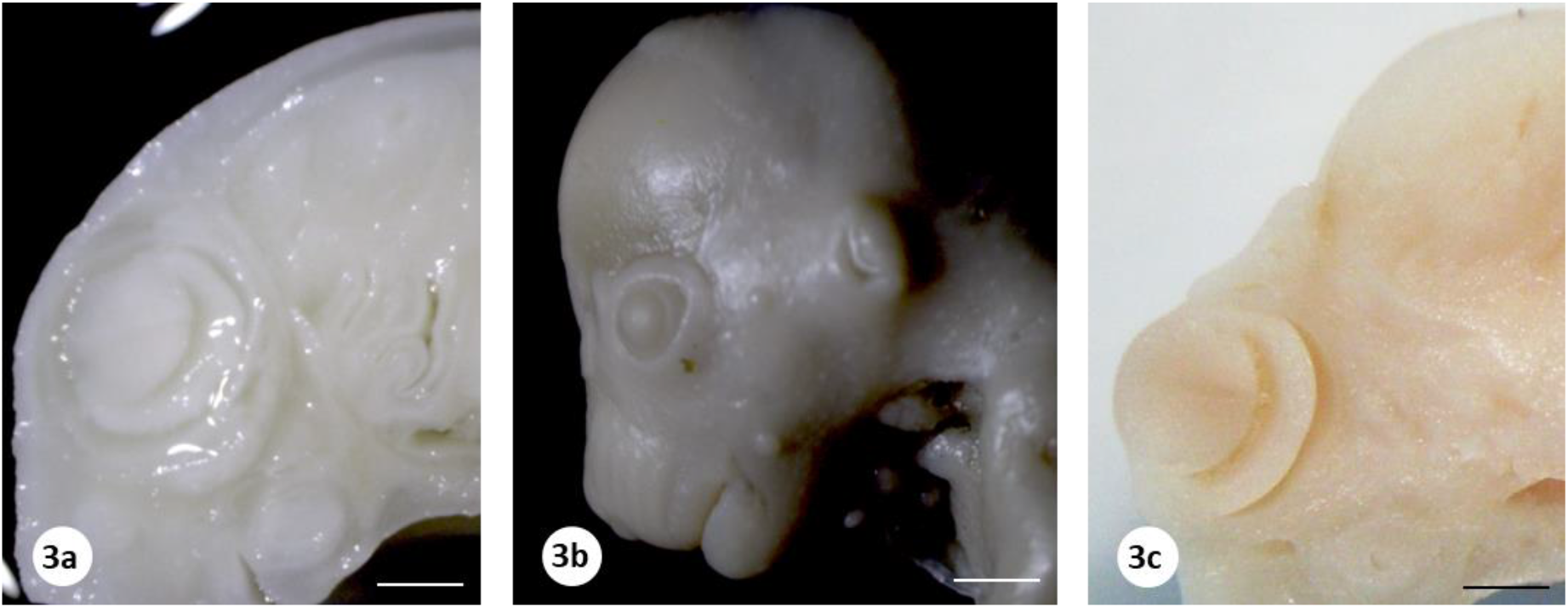
a) Photograph of a normal fetus in coronal section; b and c) CBZ-exposed foetus exhibiting open eyelid (ablepharia) and exophthalmos, respectively. Frontal view (b) and coronal section (c). Bars: 800 μm, 340μm and 750 μm, respectively

## DISCUSSION

Mammals reproduction is a complex and prolonged process that has become susceptible to exposure to external agents; the gestational period is one of the most sensitive phases. Thus, it can be considered that maternal exposure to chemical agents, can result in important effects on the embryonic-fetal organism (Damasceno et al. 2008). Both epilepsy and AEDs, such CBZ, have been associated with adverse pregnancy, fetal, and neonatal outcomes (Cassina et al. 2013; Gedzelman and Meador 2012; Kaushik et al. 2016; Li et al. 2019; Liguori and Cianfarani 2009; Thomas et al. 2017; Veiby et al. 2009; Wen et al. 2017).

However, it should be considered that the toxicokinetic of xenobiotic substances may be modified during pregnancy. In fact, women with epilepsy have an increased frequency of seizures during pregnancy due to declining serum levels of AEDs, requiring readjustment and increased doses of these drugs (Pennell 2003; Tomson 2005). This increase is mainly due to the reduction of CBZ plasma binding proteins, the presence of free CBZ in blood and the acceleration of its metabolism and elimination. The decrease in binding of CBZ to plasma proteins may lead to increased free blood fraction (Hirama et al. 2008; Tanaka, 1999).

Obstetric and perinatal complications such as increased incidence of preterm delivery, perinatal morbidity and mortality are frequent in women with epilepsy (Hirama et al. 2008). Thus, the risk-benefit of the need for a treatment with anticonvulsant drugs to control seizures should be considered. The recommendation is to continue treatment with anticonvulsant drugs during pregnancy (Czeizel and Bánhidy 2011; Weston et al. 2016). There is no consensus that anticonvulsants are safer in pregnancy, since they all have teratogenic potential. However, CBZ is still considered the recommended drug for women at risk of pregnancy. Although the CBZ is considered a human teratogen, it is an important drug option for women with epilepsy during their childbearing years because of its low cost, wide availability, relative safety for fetal outcomes and effectiveness during pregnancy (Brosh et al. 2011; Gedzelman and Meador 2012; Jentink et al. 2010; Johnson et al. 2014; Matalon et al. 2002; Pennell 2008; Tomson 2005; Vajda et al. 2013).

The female reproductive performance is a very important parameter for the analysis of the perinatal toxicity of drugs (Lemonica et al. 1996; Ortiz et al. 1979; Waynforth 1971). Maternal toxicity related to xenobiotic agents can be assessed by body weight gain during pregnancy, clinical toxicity signs or gestational alterations (Chahoud et al. 1999). Body weight is a sensitive indicator of potentially toxic chemicals in the study of reproductive toxicity (Chahoud et al. 1999; Chernoff et al. 2008; Chung et al. 2007; Kim et al. 2011).

The CBZ dose used in this study, which is equivalent to the therapeutic dose in humans, led to a reduction in body weight gain in CBZ-treated rats from DG13, evidencing that CBZ does cause maternal toxicity; nevertheless, the rats exhibited no signs of physical weakness or gestational problems. But, the change in maternal body weight, as an indicator of maternal toxicity, is not necessarily accompanied by embryofetal toxicity such as decreased fetal weight and increased malformations incidence, i.e., developmental effects are not always associated with maternal toxicity; the possibility remains that at least in some cases maternal toxicity may exacerbate the direct effects of a teratogen (Chahoud et al. 1999; Francis and Farland 1987; Hood and Rogers 2012).

It was observed in this study that CBZ interfered on several reproductive outcomes including reduction in the implantation index, average number of corpora lutea, fetal weights and crown-rump length and an increase of pre and post-implantation loss rate. A reduction in the number of corpora lutea and in the implantation index, together with an increase in the pre- and/or post-implantation loss are critical indexes for the evaluation of female fertility and should also be considered an adverse reproductive effect (Parker et al. 2012). Meanwhile, resorption rate, litter size, number of live/ dead fetuses, sex ratio, placental weight and index did not demonstrate significant alterations in CBZ-exposed pregnant rats. Pre- and post-implantation loss rates allow identify whether an environmental factor interferes with events that occurred prior to implantation or that caused changes in the post-implantation period, i.e., the maintenance of pregnancy (Roblero et al. 1987).

Pre and post-implantation loss rates allow the identification of whether an environmental factor interferes with events that occurred prior to implantation or caused changes in the post-implantation period, i.e., the maintenance of pregnancy. The rate of pre-implantation losses establishes the relation between two variables, the number of corpora lutea and the number of implantations and corresponds to the number of unfertilized oocytes and / or the embryonic losses before the endometrial implantation. An increase in pre-implantation loss may indicate an adverse effect on gamete transport, fertilization, zygote/blastula, and/or the process of implantation itself. Thus, a reduction in the implantation index is associated with an increase in the pre-implantation loss rate. CBZ may have caused alterations in the fertilization processes, implantation of blastocyst, inducing pre and/or peri-implantation losses. Changes in pre-implantation embryonic development may be caused by direct action of the xenobiotic agent on the conceptus or indirectly, due to changes in tubal secretion or modification of transport time to the uterus caused by the toxic agent, impairing the necessary timing of implantation (Roblero et al. 1987). Late embryonic-fetal viability damage is evidenced by an increase in post-implantation loss (Almeida et al. 2000).

This drug is transferred by the placenta and can accumulate in fetal tissue, inducing intrauterine growth restriction (IUGR) (Bath and Scharfman 2013; Luef 2009, Nie et al. 2016). Clinical evidences demonstrated an increase in free CBZ blood levels, further facilitating its passage through the placental barrier (Johnson et al. 2014). There is a characteristic relationship between CBZ use during pregnancy, developmental defects and fetal CBZ syndrome according to epidemiological and experimental studies (Afshar et al., 2010; Matalon et al. 2002; Wlodarczyk et al. 2012).

In developmental toxicity studies, a reduction in mean fetal body weight is usually considered to be a result of prenatal growth retardation (Chahoud and Paumgartten 2005). In this study, exposure to CBZ throughout the gestational period caused a reduction in fetal crown-rump length and body weight, along with an increase in the percentage of SPA fetuses as well as decreased rates of APA and LPA fetuses, indicating changes in intrauterine fetal growth. IUGR is the leading cause of fetal, neonatal and perinatal morbidity and mortality (Brodsky and Christou, 2004; Flenady et al. 2011; Frøen et al. 2004). The intrauterine development in mammals is the period of active cell proliferation and differentiation and it is highly sensitive to chemical insults. A number of effects, ranging from growth retardation to severe organ anomalies and functional defects, have been reported as a result of chemical exposure to embryos (Walker et al. 2000).

CBZ has effects on fetal growth and development. Experimental studies reported decreased fetal body weight and crown-rump length due to CBZ exposure, indicating IUGR and possible relation to fetal ossification retardation (Diav-Citrin et al. 2001; El-Gaafarawi and Abouel-Magd 2015; Sucheston et al. 1986).

Regarding the impact of CBZ on fetal growth and organogenesis, IUGR is inversely proportional to the CBZ dose, besides incomplete ossification of the skull, ribs and hind limbs, absence of maxilla ossification and scoliosis (El-Gaafarawi and Abouel-Magd 2015; Sucheston et al. 1986). Although it was observed a reduction in fetal weight and length due to prenatal exposure to CBZ, no significant changes were observed during the examination of skeletal anomalies. A reduction in fetal body weight may be related to changes in the ossification process, especially later ossification centers such as sternebrae V and VI of the prenatal sternum, anterior and posterior phalangeal segments of the paws, and growth retardation (Kimmel et al. 1987).

Although skeletal examination is further confused by the differences between mouse and rat, the time of laparotomy coincides with the period of peak osteogenesis. The ossification of rodent fetal bones occurs rapidly during the last 48 hours of gestation. Because of these factors, the skeletal system in rodent fetuses is immature when it is evaluated. So, reduced degree of ossification of some fetal skeleton bones does not necessarily imply that a growth retardation process has occurred. Although bone ossification advances with gestational age, an impairment of calcification of a particular bone is not necessarily secondary to a slower development of the skeleton as a whole.

The delay in prenatal ossification may be transient and does not persists in the postnatal phase (Carney and Kimmel 2007). It should also be considered that the delay in ossification depends on the age at which the fetuses were collected (Daston and Seed 2007). In addition, delayed ossification typically occurs in association with other effects on fetal growth, particularly decreased fetal weight, as found in this study. Usually, delayed ossification and decreased fetal weight are indicatives of widespread effect on fetal maturation (Daston and Seed 2007). In this study, cesarean sections were performed on the 19th day of gestation and, therefore, the skeletal changes observed in animals belonging to the CBZ-treated group should be interpreted with caution, and further studies are needed to complement this assessment.

The IUGR may be associated with oxidative stress-induced adverse intrauterine environment to the embryofetal development (Krishna and Bhalera 2011; Sharma 2016). Oxidative stress causes embryos injury due to peroxidation of membrane phospholipids and chemical modifications of different types of biomolecules (Dennery 2007).

The consequences of these damages include mitochondrial activity, embryonic development and apoptosis. Nonetheless, excessive oxidative stress is strongly associated with embryotoxicity. Over the course of the development, the delicate balance between oxidants and antioxidants can be disrupted by exogenous agents that induce reactive oxygen species (ROS) production and lead to oxidative stress. Fetal and embryonic periods are highly susceptible to tissue and organ damage by ROS. In addition, gestation itself is a physiological condition with increased metabolic demand and increased tissue oxygen requirements and, in the event of abnormalities in pregnancy, oxidative imbalance may occur, and excess free radicals can promote damage to the fetus, which already has an antioxidant defense system. It is important to emphasize that the main intracellular antioxidant component, glutathione, only reaches its maximum production at the end of gestation (Calabrese et al. 2010; Gracy et al. 2009; Wells et al. 2005).

In fact, CBZ can induce oxidative stress, resulting in excessive production of ROS and/or insufficient removal of these compounds by the antioxidant defense system. Anticonvulsant drugs may still cause direct mitochondrial damage, interfering with oxidative phosphorylation, the respiratory chain and β-oxidation, leading to decreased intracellular ATP levels, and promoting mitochondrial membrane rupture (Finsterer and Scorza 2017).

Thus, fetuses exposed to anticonvulsant drugs, including CBZ, are more susceptible to congenital malformations, such as cardiovascular, genitourinary tract and craniofacial anomalies, neural tube defects, limb anomalies, neurobehavioral alterations and IUGR (Adab et al. 2015; Brosh et al. 2011; Diav-Citrin et al. 2001; El-Sayed 1998; El-Sayed et al. 1983; Holmes 2002; Jentink et al. 2010; Jones et al. 1989; Matalon et al. 2002; Ornoy 2006; Pennell 2008; Piersma et al. 1998; Tomson et al. 2018; Vajda et al. 2013).

CBZ teratogenicity mechanisms appear to be related to the inhibition of K^+^ fast channel activation, folic acid antagonism, reduction of retinoic acid and formation of toxic metabolites such as methyl sulfonyl epoxide-CBZ (CBZE) (Amore et al. 1997; Fex et al. 1995; Raymond et al. 1995).

The teratogenicity of CBZ was investigated in Sprague-Dawley rats at doses of 0, 200, 400, and 600 mg/kg administered by gavage, from DG 7-18. The 200mg/kg dose of CBZ did not cause fetal abnormalities but reduced fetal body weight. However, the 400 and 600 mg/kg CBZ doses caused maternal toxicity evidenced by reduced body weight gain during pregnancy, reduced live fetal weight and increased visceral abnormalities. Skeletal anomalies only occurred at the highest dose. The authors suggested that CBZ is not potent at inducing malformations in rats because they only occurred at concentrations above the therapeutic range (Vorhees et al. 1990).

Afshar et al. (2010) observed that i.p. CBZ administration in pregnant mice from DG 6 to 15, at doses of 15 mg/kg/day and 30mg/kg/day, induced several external and skeletal malformations, including premature eye opening, mild to severe exophthalmos, brachygnathia, vertebral and calvarial deformities, brachydactyly, short tail and growth retardation. In this study, the occurrence of eye malformations was prevalent, mainly premature opening of both eyes. Other studies also reported congenital eye malformations (bilateral anophthalmia, bilateral microphthalmia, unilateral optic nerve coloboma) associated with the treatment of pregnant women with CBZ (Fahnehjelm et al. 1999; Kroes 2002; Sutcliffe et al. 1998). However, there was no significant alteration in the proportion of external and visceral anomalies, which could characterize the teratogenic effect. It has been reported that CBZ induces a pattern of minor anomalies (Gladstone et al. 1992).

Nonetheless, the risk associated with prenatal CBZ exposure is of considerable importance, as this medication has a variety of clinical applications. So, the prescription of CBZ must be monitored to maintain therapeutic effects for the mother but, at the same time, to minimize fetal drug exposure. In this context, human trials are limited, and experimental animal models can be used in biological research, being of great importance for medical science (Fagundes et al. 2004; Schanaider and Silva 2004). Whereas recognition drugs and other agents as potential teratogens, especially in cases where toxic effects are not as significant or where malformations occur spontaneously and frequently in the general population, experimental trials, such as this study, are essential for the risk determination in embryofetal development and may help in the interpretation of clinical studies.

## ACKNOWLEDGMENTS

The authors thank CAPES (Coordenação de Aperfeiçoamento de Pessoal de Nível Superior) and FAPESP (Fundação de Amparo à Pesquisa do Estado de São Paulo - Processo 2012/05905-9) for financial support.

## CONFLICT OF INTEREST

The authors declare that they have no conflict of interest.

## Notes

### Competing Interest Statement

The authors have declared no competing interest.

