## Supplemental data for "Carbamazepine Causes Changes in Maternal Reproductive Performance and Fetal Growth Retardation in Rats"

### Pregnant Weight

| Pregnant rat | Control Group | | | | | | | | | | | | | | | | | | | Uterus + fetuses weight |
|---|---|---|---|---|---|---|---|---|---|---|---|---|---|---|---|---|---|---|---|---|
| | Gestacional day (GD) | | | | | | | | | | | | | | | | | | | |
| | 1 | 2 | 3 | 4 | 5 | 6 | 7 | 8 | 9 | 10 | 11 | 12 | 13 | 14 | 15 | 16 | 17 | 18 | 19 | |
| 1 | 267.00 | 266.00 | 278.00 | 275.00 | 283.00 | 280.00 | 284.00 | 289.00 | 284.00 | 282.00 | 292.00 | 296.00 | 293.00 | 305.00 | 313.00 | 321.00 | 328.00 | 337.00 | 359.00 | 82.57 |
| 2 | 268.00 | 270.00 | 274.00 | 274.00 | 278.00 | 277.00 | 276.00 | 272.00 | 272.00 | 268.00 | 278.00 | 282.00 | 279.00 | 290.00 | 291.00 | 305.00 | 316.00 | 333.00 | 338.00 | 77.74 |
| 3 | 241.00 | 244.00 | 248.00 | 251.00 | 244.00 | 245.00 | 254.00 | 249.00 | 256.00 | 255.00 | 258.00 | 259.00 | 266.00 | 274.00 | 274.00 | 282.00 | 289.00 | 304.00 | 307.00 | 70.61 |
| 4 | 244.00 | 251.00 | 245.00 | 240.00 | 253.00 | 255.00 | 257.00 | 260.00 | 260.00 | 259.00 | 266.00 | 271.00 | 269.00 | 288.00 | 289.00 | 296.00 | 300.00 | 317.00 | 324.00 | 74.52 |
| 5 | 244.00 | 251.00 | 256.00 | 258.00 | 252.00 | 258.00 | 249.00 | 252.00 | 259.00 | 263.00 | 263.00 | 271.00 | 272.00 | 269.00 | 283.00 | 291.00 | 297.00 | 308.00 | 322.00 | 74.06 |
| 6 | 282.00 | 282.00 | 286.00 | 284.00 | 282.00 | 292.00 | 286.00 | 293.00 | 297.00 | 306.00 | 303.00 | 304.00 | 314.00 | 310.00 | 319.00 | 328.00 | 340.00 | 347.00 | 364.00 | 83.72 |
| 7 | 275.00 | 274.00 | 272.00 | 281.00 | 275.00 | 280.00 | 280.00 | 284.00 | 285.00 | 289.00 | 295.00 | 295.00 | 304.00 | 308.00 | 313.00 | 319.00 | 321.00 | 335.00 | 359.00 | 82.57 |
| 8 | 243.00 | 244.00 | 250.00 | 249.00 | 254.00 | 252.00 | 256.00 | 252.00 | 260.00 | 267.00 | 259.00 | 279.00 | 279.00 | 283.00 | 285.00 | 292.00 | 306.00 | 312.00 | 328.00 | 75.44 |
| 9 | 255.00 | 248.00 | 253.00 | 251.00 | 258.00 | 259.00 | 257.00 | 257.00 | 269.00 | 272.00 | 277.00 | 284.00 | 287.00 | 296.00 | 301.00 | 314.00 | 323.00 | 337.00 | 348.00 | 80.04 |
| **Mean** | **257.67** | **258.89** | **262.44** | **262.56** | **264.33** | **266.44** | **266.56** | **267.56** | **271.33** | **273.44** | **276.78** | **282.33** | **284.78** | **291.44** | **296.44** | **305.33** | **313.33** | **325.56** | **338.78** | **77.92** |
| **SD** | **15.65** | **14.24** | **15.08** | **16.07** | **15.01** | **16.04** | **14.63** | **17.30** | **14.39** | **16.23** | **16.69** | **14.28** | **16.31** | **14.65** | **15.73** | **15.95** | **16.49** | **15.40** | **19.89** | **4.58** |

| Pregnant rat | CBZ Group | | | | | | | | | | | | | | | | | | | Uterus + fetuses weight |
|---|---|---|---|---|---|---|---|---|---|---|---|---|---|---|---|---|---|---|---|---|
| | Gestacional day (GD) | | | | | | | | | | | | | | | | | | | |
| | 1 | 2 | 3 | 4 | 5 | 6 | 7 | 8 | 9 | 10 | 11 | 12 | 13 | 14 | 15 | 16 | 17 | 18 | 19 | |
| 1 | 251.00 | 253.00 | 258.00 | 257.00 | 260.00 | 258.00 | 261.00 | 260.00 | 268.00 | 270.00 | 272.00 | 267.00 | 273.00 | 273.00 | 286.00 | 282.00 | 293.00 | 301.00 | 307.00 | 61.40 |
| 2 | 235.00 | 228.00 | 239.00 | 235.00 | 238.00 | 237.00 | 238.00 | 240.00 | 245.00 | 247.00 | 250.00 | 247.00 | 255.00 | 257.00 | 260.00 | 263.00 | 266.00 | 276.00 | 294.00 | 58.80 |
| 3 | 274.00 | 264.00 | 274.00 | 276.00 | 275.00 | 278.00 | 278.00 | 284.00 | 283.00 | 291.00 | 294.00 | 297.00 | 294.00 | 299.00 | 302.00 | 299.00 | 313.00 | 325.00 | 346.00 | 69.20 |
| 4 | 253.00 | 259.00 | 263.00 | 263.00 | 266.00 | 271.00 | 266.00 | 270.00 | 280.00 | 272.00 | 279.00 | 277.00 | 287.00 | 285.00 | 283.00 | 295.00 | 298.00 | 310.00 | 315.00 | 63.00 |
| 5 | 250.00 | 250.00 | 257.00 | 256.00 | 260.00 | 263.00 | 261.00 | 261.00 | 269.00 | 264.00 | 268.00 | 266.00 | 267.00 | 268.00 | 270.00 | 277.00 | 289.00 | 298.00 | 315.00 | 63.00 |
| 6 | 239.00 | 238.00 | 246.00 | 248.00 | 250.00 | 251.00 | 249.00 | 254.00 | 260.00 | 258.00 | 266.00 | 262.00 | 268.00 | 269.00 | 275.00 | 283.00 | 284.00 | 307.00 | 308.00 | 61.60 |
| 7 | 257.00 | 263.00 | 264.00 | 267.00 | 273.00 | 274.00 | 274.00 | 272.00 | 268.00 | 271.00 | 276.00 | 277.00 | 278.00 | 274.00 | 278.00 | 283.00 | 280.00 | 285.00 | 295.00 | 59.00 |
| 8 | 263.00 | 263.00 | 272.00 | 277.00 | 277.00 | 273.00 | 272.00 | 276.00 | 273.00 | 275.00 | 264.00 | 263.00 | 268.00 | 266.00 | 266.00 | 263.00 | 275.00 | 296.00 | 313.00 | 62.60 |
| 9 | 277.00 | 278.00 | 283.00 | 288.00 | 293.00 | 288.00 | 293.00 | 288.00 | 290.00 | 291.00 | 292.00 | 299.00 | 294.00 | 303.00 | 311.00 | 313.00 | 310.00 | 325.00 | 339.00 | 67.80 |
| 10 | 258.00 | 261.00 | 261.00 | 267.00 | 266.00 | 266.00 | 266.00 | 263.00 | 261.00 | 271.00 | 267.00 | 273.00 | 272.00 | 276.00 | 284.00 | 279.00 | 288.00 | 295.00 | 295.00 | 59.00 |
| **Mean** | **255.70** | **255.70** | **261.70** | **263.40** | **265.80** | **265.90** | **265.80** | **266.80** | **269.70** | **271.00** | **272.80** | **272.80** | **275.60** | **277.00** | **281.50** | **283.70** | **289.60** | **301.80** | **312.70** | **62.54** |
| **SD** | **13.41** | **14.27** | **12.98** | **15.33** | **15.24** | **14.55** | **15.26** | **14.31** | **12.88** | **13.38** | **13.20** | **15.87** | **12.68** | **14.59** | **15.66** | **15.45** | **14.69** | **15.68** | **17.77** | **3.55** |

### Maternal Reproductive Outcome

| Animals | Implantation sites | | | Resorption | | | *Corpora lutea* | | | Pre-implantation loss rate | Post-implantation loss rate | Resorption rate | Fetus | | | | | Implantation index | Sex ratio | Fetus | | Placental weight (g) | Placental index |
|---|---|---|---|---|---|---|---|---|---|---|---|---|---|---|---|---|---|---|---|---|---|---|---|
| | R | L | T | R | L | T | R | L | T | | | | Live | Dead | T | F | M | | | Weight (g) | Crown-rump length (mm) | | |
| Control Group | | | | | | | | | | | | | | | | | | | | | | | |
| 1 | 8.00 | 5.00 | 13.00 | 0.00 | 0.00 | 0.00 | 8.00 | 5.00 | 13.00 | 0.00 | 0.00 | 0.00 | 13.00 | 0.00 | 13.00 | 7.00 | 6.00 | 100.00 | 0.86 | 1.44 | 23.23 | 0.33 | 0.23 |
| 2 | 7.00 | 7.00 | 14.00 | 0.00 | 0.00 | 0.00 | 7.00 | 7.00 | 14.00 | 0.00 | 7.00 | 0.00 | 13.00 | 1.00 | 14.00 | 6.00 | 8.00 | 100.00 | 1.33 | 2.19 | 27.71 | 0.39 | 0.18 |
| 3 | 9.00 | 3.00 | 12.00 | 0.00 | 0.00 | 0.00 | 9.00 | 3.00 | 12.00 | 0.00 | 0.00 | 0.00 | 12.00 | 0.00 | 12.00 | 7.00 | 5.00 | 100.00 | 0.71 | 1.60 | 24.42 | 0.39 | 0.24 |
| 4 | 4.00 | 6.00 | 10.00 | 0.00 | 1.00 | 1.00 | 5.00 | 6.00 | 11.00 | 9.00 | 10.00 | 10.00 | 9.00 | 0.00 | 9.00 | 5.00 | 4.00 | 90.91 | 0.80 | 1.59 | 23.67 | 0.47 | 0.30 |
| 5 | 4.00 | 11.00 | 15.00 | 0.00 | 0.00 | 0.00 | 5.00 | 10.00 | 15.00 | 0.00 | 0.00 | 0.00 | 15.00 | 0.00 | 15.00 | 10.00 | 5.00 | 100.00 | 0.50 | 1.49 | 23.10 | 0.42 | 0.28 |
| 6 | 7.00 | 5.00 | 12.00 | 0.00 | 1.00 | 1.00 | 7.00 | 5.00 | 12.00 | 0.00 | 8.00 | 8.33 | 11.00 | 0.00 | 11.00 | 5.00 | 6.00 | 100.00 | 1.20 | 1.56 | 23.72 | 0.41 | 0.26 |
| 7 | 6.00 | 7.00 | 13.00 | 0.00 | 1.00 | 1.00 | 6.00 | 7.00 | 13.00 | 0.00 | 8.00 | 7.69 | 12.00 | 0.00 | 12.00 | 6.00 | 6.00 | 100.00 | 1.00 | 1.56 | 23.82 | 0.46 | 0.29 |
| 8 | 5.00 | 6.00 | 11.00 | 0.00 | 0.00 | 0.00 | 5.00 | 6.00 | 11.00 | 0.00 | 0.00 | 0.00 | 11.00 | 0.00 | 11.00 | 6.00 | 5.00 | 100.00 | 0.83 | 1.52 | 24.18 | 0.47 | 0.31 |
| 9 | 7.00 | 5.00 | 12.00 | 1.00 | 0.00 | 1.00 | 7.00 | 5.00 | 12.00 | 0.00 | 8.33 | 8.33 | 11.00 | 0.00 | 11.00 | 4.00 | 7.00 | 100.00 | 1.75 | 1.64 | 24.09 | 0.41 | 0.25 |
| **Sun** | **57.00** | **55.00** | **112.00** | **1.00** | **3.00** | **4.00** | **59.00** | **54.00** | **113.00** | | | | **107.00** | **1.00** | **108.00** | **56.00** | **52.00** | | | | | | |
| **Mean** | | | **12.44** | | | **0.44** | | | **12.56** | **1.00** | **4.59** | **3.82** | **11.89** | **0.11** | **12.00** | **6.22** | **5.78** | **98.99** | **1.00** | **1.62** | **24.22** | **0.42** | **0.26** |
| **DP** | | | **1.51** | | | **0.53** | | | **1.33** | **3.00** | **4.42** | **4.57** | **1.69** | **0.33** | **1.80** | **1.72** | **1.20** | **3.03** | **0.38** | **0.22** | **1.38** | **0.05** | **0.04** |
| **Median** | | | **12.00** | | | **0.00** | | | **12.00** | **0.00** | **7.00** | **0.00** | **12.00** | **0.00** | **12.00** | **6.00** | **6.00** | **100.00** | **0.86** | **1.56** | **23.82** | **0.41** | **0.26** |
| **IQ1** | | | **12.00** | | | **0.00** | | | **12.00** | **0.00** | **0.00** | **0.00** | **11.00** | **0.00** | **11.00** | **5.00** | **5.00** | **100.00** | **0.80** | **1.52** | **23.67** | **0.39** | **0.24** |
| **IQ3** | | | **13.00** | | | **1.00** | | | **13.00** | **0.00** | **8.00** | **8.33** | **13.00** | **0.00** | **13.00** | **7.00** | **6.00** | **100.00** | **1.20** | **1.60** | **24.18** | **0.46** | **0.29** |
| CBZ Group | | | | | | | | | | | | | | | | | | | | | | | |
| 1 | 4.00 | 7.00 | 11.00 | 1.00 | 1.00 | 2.00 | 9.00 | 11.00 | 20.00 | 45.00 | 18.00 | 18.00 | 9.00 | 0.00 | 9.00 | 5.00 | 4.00 | 55.00 | 0.80 | 1.17 | 22.20 | 0.39 | 0.33 |
| 2 | 8.00 | 5.00 | 13.00 | 1.00 | 0.00 | 1.00 | 9.00 | 4.00 | 13.00 | 0.00 | 69.00 | 8.00 | 4.00 | 8.00 | 12.00 | 6.00 | 6.00 | 100.00 | 1.00 | 1.30 | 22.17 | 0.33 | 0.25 |
| 3 | 6.00 | 7.00 | 13.00 | 0.00 | 1.00 | 1.00 | 7.00 | 7.00 | 14.00 | 7.00 | 8.00 | 8.00 | 12.00 | 0.00 | 12.00 | 8.00 | 4.00 | 92.86 | 0.50 | 1.51 | 24.58 | 0.42 | 0.28 |
| 4 | 1.00 | 8.00 | 9.00 | 1.00 | 1.00 | 2.00 | 8.00 | 6.00 | 14.00 | 36.00 | 22.00 | 22.00 | 7.00 | 0.00 | 7.00 | 3.00 | 4.00 | 64.29 | 1.33 | 1.71 | 24.43 | 0.42 | 0.25 |
| 5 | 8.00 | 5.00 | 13.00 | 1.00 | 0.00 | 1.00 | 8.00 | 5.00 | 13.00 | 0.00 | 8.00 | 8.00 | 12.00 | 0.00 | 12.00 | 5.00 | 7.00 | 100.00 | 1.40 | 1.38 | 23.18 | 0.35 | 0.25 |
| 6 | 5.00 | 7.00 | 12.00 | 1.00 | 0.00 | 1.00 | 5.00 | 7.00 | 12.00 | 0.00 | 8.00 | 8.00 | 11.00 | 0.00 | 11.00 | 7.00 | 4.00 | 100.00 | 0.57 | 1.54 | 23.46 | 0.38 | 0.25 |
| 7 | 9.00 | 3.00 | 12.00 | 1.00 | 0.00 | 1.00 | 9.00 | 5.00 | 14.00 | 14.00 | 8.00 | 8.00 | 11.00 | 0.00 | 11.00 | 5.00 | 6.00 | 85.71 | 1.20 | 1.18 | 21.36 | 0.32 | 0.27 |
| 8 | 6.00 | 10.00 | 16.00 | 0.00 | 0.00 | 0.00 | 6.00 | 10.00 | 16.00 | 0.00 | 0.00 | 0.00 | 16.00 | 0.00 | 16.00 | 9.00 | 7.00 | 100.00 | 0.78 | 1.27 | 22.38 | 0.39 | 0.31 |
| 9 | 10.00 | 3.00 | 13.00 | 1.00 | 1.00 | 1.00 | 12.00 | 3.00 | 15.00 | 13.00 | 15.00 | 8.00 | 11.00 | 0.00 | 11.00 | 5.00 | 6.00 | 86.67 | 1.20 | 1.42 | 22.91 | 0.43 | 0.30 |
| 10 | 1.00 | 2.00 | 3.00 | 0.00 | 0.00 | 0.00 | 8.00 | 5.00 | 13.00 | 77.00 | 0.00 | 0.00 | 3.00 | 0.00 | 3.00 | 3.00 | 0.00 | 23.08 | 0.00 | 1.38 | 23.00 | 0.57 | 0.41 |
| **Sun** | **58.00** | **57.00** | **115.00** | **7.00** | **4.00** | **10.00** | **81.00** | **63.00** | **144.00** | | | | **96.00** | **8.00** | **104.00** | **56.00** | **48.00** | | | | | | **2.32** |
| **Mean** | | | **11.50** | | | **1.00** | | | **14.40** | **19.20** | **15.60** | **8.80** | **9.60** | **0.80** | **10.40** | **5.60** | **4.80** | **81.58** | **0.87** | **1.42** | **23.16** | **0.41** | **0.29** |
| **DP** | | | **3.47** | | | **0.67** | | | **2.27** | **25.77** | **20.05** | **6.81** | **3.95** | **2.53** | **3.47** | **1.96** | **2.10** | **26.48** | **0.49** | **0.16** | **1.05** | **0.07** | **0.06** |
| **Median** | | | **12.50** | | | **1.00** | | | **14.00** | **10.00** | **8.00** | **8.00** | **11.00** | **0.00** | **11.00** | **5.00** | **5.00** | **89.76** | **0.99** | **1.40** | **23.09** | **0.41** | **0.27** |
| **IQ1** | | | **11.25** | | | **1.00** | | | **13.00** | **0.00** | **8.00** | **8.00** | **7.50** | **0.00** | **9.50** | **5.00** | **4.00** | **80.36** | **0.55** | **1.35** | **22.78** | **0.37** | **0.25** |
| **IQ3** | | | **13.00** | | | **1.00** | | | **14.75** | **30.50** | **17.25** | **8.00** | **11.75** | **0.00** | **12.00** | **6.75** | **6.00** | **100.00** | **1.23** | **1.52** | **23.70** | **0.42** | **0.30** |

### Fetal Outcome

### Control Group

| Animals | Fetus | | | | | | | | | | Placental weight (g) | | |
|---|---|---|---|---|---|---|---|---|---|---|---|---|---|
| | R | | | | L | | | | Mean | | R | L | Placental weight mean |
| | Weight (g) | Weight classification | Crown-rump length (mm) | Sex | Weight (g) | Weight classification | Crown-rump length (mm) | Sex | Weight (g) | Crown-rump length (mm) | | | |
| 1 | 1.51 | APA | 24 | m | 1.48 | APA | 22 | m | 1.44 | 23.23 | 0.34 | 0.36 | 0.33 |
| | 1.39 | APA | 23 | m | 1.31 | APA | 21 | m | | | 0.27 | 0.29 | |
| | 1.36 | APA | 23 | m | 1.47 | APA | 25 | f | | | 0.3 | 0.32 | |
| | 1.51 | APA | 24 | f | 1.48 | APA | 23 | f | | | 0.35 | 0.34 | |
| | 1.42 | APA | 23 | f | 1.47 | APA | 23 | f | | | 0.42 | 0.31 | |
| | 1.5 | APA | 24 | m | | | | | | | 0.38 | | |
| | 1.39 | APA | 23 | f | | | | | | | 0.31 | | |
| | 1.44 | APA | 24 | f | | | | | | | 0.31 | | |
| 2 | 2.31 | LPA | 30 | f | 2.33 | LPA | 31 | m | 2.19 | 27.71 | 0.43 | 0.42 | 0.39 |
| | 2.25 | LPA | 28 | f | 2.08 | LPA | 29 | m | | | 0.46 | 0.48 | |
| | 2.3 | LPA | 29 | m | 2.15 | LPA | 29 | m | | | 0.38 | 0.38 | |
| | 1.3 | APA | 16 | m | 2.35 | LPA | 30 | f | | | 0.17 | 0.39 | |
| | 2.57 | LPA | 27 | m | 2.14 | LPA | 29 | f | | | 0.35 | 0.36 | |
| | 2.26 | LPA | 29 | m | 2.24 | LPA | 28 | f | | | 0.44 | 0.4 | |
| | 2.21 | LPA | 25 | f | 2.27 | LPA | 28 | m | | | 0.4 | 0.35 | |
| 3 | 1.53 | APA | 23 | f | 1.56 | APA | 25 | f | 1.6 | 24.42 | 0.37 | 0.46 | 0.39 |
| | 1.64 | APA | 25 | f | 1.72 | APA | 25 | f | | | 0.4 | 0.48 | |
| | 1.55 | APA | 23 | m | 1.71 | APA | 24 | m | | | 0.31 | 0.51 | |
| | 1.69 | APA | 26 | m | | | | | | | 0.39 | | |
| | 1.5 | APA | 24 | f | | | | | | | 0.32 | | |
| | 1.54 | APA | 25 | f | | | | | | | 0.3 | | |
| | 1.66 | APA | 25 | m | | | | | | | 0.34 | | |
| | 1.67 | APA | 25 | m | | | | | | | 0.44 | | |
| | 1.48 | APA | 23 | f | | | | | | | 0.33 | | |
| 4 | 1.58 | APA | 23 | m | 1.62 | APA | 26 | f | 1.59 | 23.67 | 0.43 | 0.59 | 0.47 |
| | 1.68 | APA | 22 | f | 1.59 | APA | 23 | f | | | 0.47 | 0.43 | |
| | 1.66 | APA | 25 | f | 1.49 | APA | 23 | m | | | 0.44 | 0.4 | |
| | 1.72 | APA | 24 | m | 1.52 | APA | 24 | m | | | 0.46 | 0.52 | |
| | | | | | 1.42 | APA | 23 | f | | | | 0.49 | |
| 5 | 1.24 | SPA | 23 | m | 1.1 | SPA | 21 | f | 1.49 | 23.1 | 0.36 | 0.41 | 0.42 |
| | 1.72 | APA | 25 | f | 1.52 | APA | 23 | f | | | 0.47 | 0.42 | |
| | 1.62 | APA | 23 | f | 1.44 | APA | 22 | m | | | 0.47 | 0.4 | |
| | 1.75 | APA | 26 | f | 1.55 | APA | 22 | f | | | 0.55 | 0.39 | |
| | | | | | 1.48 | APA | 22 | f | | | | 0.43 | |
| | | | | | 1.39 | APA | 23 | m | | | | 0.38 | |
| | | | | | 1.41 | APA | 23 | m | | | | 0.45 | |
| | | | | | 1.5 | APA | 23 | f | | | | 0.4 | |
| | | | | | 1.51 | APA | 24 | f | | | | 0.43 | |
| | | | | | 1.51 | APA | 22 | m | | | | 0.37 | |
| | | | | | 1.58 | APA | 24 | f | | | | 0.35 | |
| 6 | 1.46 | APA | 23 | m | 1.54 | APA | 24 | f | 1.56 | 23.72 | 0.45 | 0.41 | 0.41 |
| | 1.54 | APA | 25 | f | 1.58 | APA | 25 | m | | | 0.35 | 0.53 | |
| | 1.49 | APA | 23 | m | 1.63 | APA | 25 | m | | | 0.37 | 0.48 | |
| | 1.76 | APA | 23 | m | 1.45 | APA | 23 | f | | | 0.48 | 0.33 | |
| | 1.58 | APA | 23 | m | | | | | | | 0.39 | | |
| | 1.6 | APA | 24 | f | | | | | | | 0.39 | | |
| | 1.56 | APA | 23 | f | | | | | | | 0.39 | | |
| 7 | 1.55 | APA | 24 | m | 1.47 | APA | 24 | f | 1.56 | 23.82 | 0.41 | 0.48 | 0.46 |
| | 1.64 | APA | 25 | m | 1.51 | APA | 23 | f | | | 0.42 | 0.44 | |
| | 1.57 | APA | 24 | f | 1.7 | APA | 23 | m | | | 0.46 | 0.52 | |
| | 1.72 | APA | 24 | m | 1.54 | APA | 22 | f | | | 0.59 | 0.39 | |
| | 1.63 | APA | 23 | m | 1.57 | APA | 23 | m | | | 0.47 | 0.39 | |
| | 1.6 | APA | 23 | f | 1.56 | APA | 24 | f | | | 0.42 | 0.48 | |
| 8 | 1.45 | APA | 23 | f | 1.54 | APA | 25 | m | 1.52 | 24.18 | 0.39 | 0.49 | 0.47 |
| | 1.6 | APA | 25 | f | 1.58 | APA | 25 | f | | | 0.44 | 0.52 | |
| | 1.61 | APA | 25 | m | 1.61 | APA | 25 | f | | | 0.45 | 0.56 | |
| | 1.55 | APA | 25 | m | 1.54 | APA | 24 | f | | | 0.41 | 0.47 | |
| | 1.64 | APA | 25 | f | 1.09 | SPA | 20 | m | | | 0.47 | 0.46 | |
| | | | | | 1.55 | APA | 24 | f | | | | 0.48 | |
| 9 | 1.58 | APA | 24 | m | 1.47 | APA | 23 | f | 1.64 | 24.09 | 0.39 | 0.35 | 0.41 |
| | 1.7 | APA | 25 | m | 1.59 | APA | 23 | m | | | 0.39 | 0.5 | |
| | 1.6 | APA | 24 | m | 1.6 | APA | 23 | f | | | 0.44 | 0.41 | |
| | 1.88 | APA | 26 | f | 1.67 | APA | 25 | m | | | 0.36 | 0.39 | |
| | 1.6 | APA | 23 | f | 1.7 | APA | 25 | m | | | 0.44 | 0.42 | |
| | 1.64 | APA | 24 | m | | | | | | | 0.44 | | |

CBZ Group

| Animals | Fetus | | | | | | | | | | Placental weight (g) | | |
|---|---|---|---|---|---|---|---|---|---|---|---|---|---|
| | R | | | | L | | | | Mean | | R | L | Placental weight mean |
| | Weight (g) | Weight classification | Crown-rump length (mm) | Sex | Weight (g) | Weight classification | Crown-rump length (mm) | Sex | Weight (g) | Crown-rump length (mm) | | | |
| 1 | 1.27 | APA | 22.00 | m | 1.29 | APA | 24.00 | m | 1.17 | 22.20 | 0.47 | 0.41 | 0.39 |
| | 1.13 | SPA | 23.00 | f | 1.37 | APA | 25.00 | f | | | 0.28 | 0.37 | |
| | 1.35 | APA | 23.00 | f | 1.35 | APA | 23.00 | f | | | 0.46 | 0.36 | |
| | | | | | 0.73 | SPA | 17.00 | m | | | | 0.39 | |
| | | | | | 1.20 | SPA | 23.00 | f | | | | 0.44 | |
| | | | | | 0.90 | SPA | 20.00 | m | | | | 0.39 | |
| 2 | 1.18 | SPA | 22.00 | f | 1.35 | APA | 22.00 | m | 1.30 | 22.17 | 0.31 | 0.36 | 0.33 |
| | 1.10 | SPA | 23.00 | m | 1.31 | APA | 22.00 | m | | | 0.37 | 0.37 | |
| | 1.30 | APA | 21.00 | m | 1.49 | APA | 25.00 | f | | | 0.31 | 0.41 | |
| | 1.24 | SPA | 22.00 | m | | | | | | | 0.34 | | |
| | 1.35 | APA | 23.00 | f | | | | | | | 0.31 | | |
| | 1.42 | APA | 21.00 | f | | | | | | | 0.36 | | |
| | 1.29 | APA | 22.00 | f | | | | | | | 0.34 | | |
| | 1.26 | APA | 22.00 | m | | | | | | | 0.32 | | |
| | 1.36 | APA | 21.00 | f | | | | | | | 0.25 | | |
| 3 | 1.55 | APA | 25.00 | m | 1.43 | APA | 26.00 | f | 1.51 | 24.58 | 0.44 | 0.45 | 0.42 |
| | 1.40 | APA | 24.00 | m | 1.56 | APA | 24.00 | f | | | 0.50 | 0.40 | |
| | 1.52 | APA | 26.00 | m | 1.62 | APA | 26.00 | m | | | 0.41 | 0.39 | |
| | 1.49 | APA | 24.00 | f | 1.47 | APA | 24.00 | f | | | 0.48 | 0.36 | |
| | 1.47 | APA | 24.00 | f | 1.64 | APA | 24.00 | f | | | 0.35 | 0.44 | |
| | 1.55 | APA | 25.00 | f | 1.47 | APA | 23.00 | f | | | 0.39 | 0.43 | |
| 4 | | | | | 1.70 | APA | 25.00 | f | 1.71 | 24.43 | | 0.50 | 0.42 |
| | | | | | 1.72 | APA | 24.00 | m | | | | 0.35 | |
| | | | | | 2.02 | LPA | 25.00 | f | | | | 0.39 | |
| | | | | | 1.71 | APA | 25.00 | m | | | | 0.44 | |
| | | | | | 1.67 | APA | 25.00 | m | | | | 0.41 | |
| | | | | | 1.64 | APA | 24.00 | m | | | | 0.39 | |
| | | | | | 1.57 | APA | 23.00 | f | | | | 0.52 | |
| 5 | 1.23 | SPA | 22.00 | f | 1.12 | SPA | 22.00 | f | 1.38 | 23.18 | 0.27 | 0.27 | 0.35 |
| | 1.34 | APA | 22.00 | m | 1.38 | APA | 22.00 | m | | | 0.33 | 0.27 | |
| | 1.50 | APA | 24.00 | m | 1.40 | APA | 24.00 | f | | | 0.43 | 0.35 | |
| | 1.37 | APA | 23.00 | f | 1.50 | APA | 23.00 | m | | | 0.28 | 0.33 | |
| | 1.50 | APA | 24.00 | m | 1.51 | APA | 24.00 | m | | | 0.42 | 0.43 | |
| | 1.30 | APA | 22.00 | m | | | | | | | 0.37 | | |
| | 1.41 | APA | 25.00 | f | | | | | | | 0.47 | | |
| 6 | 1.66 | APA | 24.00 | m | 1.54 | APA | 24.00 | f | 1.54 | 23.46 | 0.31 | 0.34 | 0.38 |
| | 1.56 | APA | 25.00 | m | 1.48 | APA | 22.00 | f | | | 0.42 | 0.37 | |
| | 1.47 | APA | 23.00 | f | 1.68 | APA | 25.00 | m | | | 0.36 | 0.49 | |
| | 1.62 | APA | 24.00 | f | 1.58 | APA | 23.00 | m | | | 0.44 | 0.35 | |
| | | | | | 1.42 | APA | 23.00 | f | | | | 0.35 | |
| | | | | | 1.37 | APA | 22.00 | f | | | | 0.40 | |
| | | | | | 1.57 | APA | 23.00 | f | | | | 0.39 | |
| 7 | 1.31 | APA | 23.00 | f | 0.49 | SPA | 19.00 | f | 1.18 | 21.36 | 0.39 | 0.19 | 0.32 |
| | 1.31 | APA | 23.00 | f | 1.27 | APA | 22.00 | m | | | 0.35 | 0.30 | |
| | 1.28 | APA | 22.00 | m | 1.20 | SPA | 21.00 | m | | | 0.39 | 0.27 | |
| | 1.27 | APA | 22.00 | f | | | | | | | 0.30 | | |
| | 1.34 | APA | 21.00 | m | | | | | | | 0.36 | | |
| | 1.21 | SPA | 21.00 | m | | | | | | | 0.31 | | |
| | 1.12 | SPA | 20.00 | f | | | | | | | 0.39 | | |
| | 1.19 | SPA | 21.00 | m | | | | | | | 0.31 | | |
| 8 | 1.32 | APA | 23.00 | f | 1.06 | SPA | 21.00 | f | 1.27 | 22.38 | 0.38 | 0.44 | 0.39 |
| | 1.17 | SPA | 22.00 | f | 1.31 | APA | 22.00 | m | | | 0.35 | 0.39 | |
| | 1.36 | APA | 23.00 | f | 1.37 | APA | 23.00 | m | | | 0.42 | 0.42 | |
| | 1.35 | APA | 23.00 | f | 1.25 | SPA | 23.00 | f | | | 0.38 | 0.43 | |
| | 1.49 | APA | 25.00 | m | 1.33 | APA | 23.00 | f | | | 0.39 | 0.50 | |
| | 1.30 | APA | 23.00 | m | 1.10 | SPA | 21.00 | f | | | 0.31 | 0.31 | |
| | | | | | 1.26 | APA | 21.00 | m | | | | 0.37 | |
| | | | | | 1.31 | APA | 22.00 | m | | | | 0.46 | |
| | | | | | 1.27 | APA | 22.00 | m | | | | 0.34 | |
| | | | | | 1.10 | SPA | 21.00 | f | | | | 0.40 | |
| 9 | 1.39 | APA | 23.00 | f | 1.47 | APA | 25.00 | m | 1.42 | 22.91 | 0.42 | 0.54 | 0.43 |
| | 1.33 | APA | 22.00 | m | 1.33 | APA | 23.00 | m | | | 0.52 | 0.39 | |
| | 1.48 | APA | 23.00 | f | | | | | | | 0.44 | | |
| | 1.39 | APA | 23.00 | f | | | | | | | 0.39 | | |
| | 1.34 | APA | 22.00 | m | | | | | | | 0.35 | | |
| | 1.52 | APA | 23.00 | f | | | | | | | 0.48 | | |
| | 1.54 | APA | 23.00 | m | | | | | | | 0.44 | | |
| | 1.51 | APA | 23.00 | f | | | | | | | 0.50 | | |
| | 1.35 | APA | 22.00 | m | | | | | | | 0.35 | | |
| 10 | 1.51 | APA | 24.00 | f | 1.44 | APA | 24.00 | f | 1.38 | 23.00 | 0.68 | 0.52 | 0.57 |
| | | | | | 1.19 | SPA | 21.00 | f | | | | 0.52 | |

**Fetal examination**

| | External anomalies | | Visceral anomalies | | Skeletal anomalies | |
|---|---|---|---|---|---|---|
| **Control Group** | | | | | | |
| Litter | Fetus total | Anomalies types | Fetus total | Anomalies types | Fetus total | Anomalies types |
| 1 | 13 | 0 | 7 | 0 | 6 | 0 |
| 2 | 14 | 0 | 7 | 0 | 7 | 0 |
| 3 | 12 | 0 | 6 | 0 | 6 | 0 |
| 4 | 9 | 0 | 4 | 0 | 5 | 0 |
| 5 | 15 | 0 | 7 | 0 | 8 | 0 |
| 6 | 11 | 0 | 6 | 0 | 5 | 0 |
| 7 | 12 | 0 | 6 | 0 | 6 | 0 |
| 8 | 11 | 0 | 6 | 0 | 5 | 0 |
| 9 | 11 | 0 | 5 | 0 | 6 | 0 |
| **Total** | **108** | **0** | **54** | **0** | **54** | **0** |
| **CBZ Group** | | | | | | |
| Litter | Fetus total | Anomalies types | Fetus total | Anomalies types | Fetus total | Anomalies types |
| 1 | 9 | Ablepharia 1 | 5 | 0 | 4 | 0 |
| 2 | 12 | Ablepharia 1<br>Gastroschisis 1 | 6 | 0 | 6 | 0 |
| 3 | 12 | 0 | 6 | 0 | 6 | 0 |
| 4 | 7 | 0 | 3 | 0 | 4 | 0 |
| 5 | 12 | Ablepharia 1 | 6 | 0 | 6 | 0 |
| 6 | 11 | 0 | 5 | 0 | 6 | 0 |
| 7 | 11 | 0 | 6 | 0 | 5 | Unossified hindlimbs 1 |
| 8 | 16 | 0 | 8 | Ectopic testis 1 | 8 | 0 |
| 9 | 11 | 0 | 5 | 0 | 6 | 0 |
| 10 | 3 | Exophthalmos bilateral 1 | 2 | Macrophthalmia 1 | 1 | 0 |
| **Total** | **104** | **5** | **52** | **2** | **52** | **1** |
